# Older adults sacrifice response speed to preserve multisensory integration performance

**DOI:** 10.1101/474882

**Authors:** Samuel A. Jones, Ulrik Beierholm, David Meijer, Uta Noppeney

## Abstract

Ageing has been shown to impact multisensory perception, but the underlying computational mechanisms are unclear. For effective interactions with the environment, observers should integrate signals that share a common source, weighted by their reliabilities, and segregate those from separate sources. Observers are thought to accumulate evidence about the world’s causal structure over time until a decisional threshold is reached.

Combining psychophysics and Bayesian modelling, we investigated how ageing affects audiovisual perception of spatial signals. Older and younger adults were comparable in their final localisation and common-source judgement responses under both speeded and unspeeded conditions, but were disproportionately slower for audiovisually incongruent trials.

Bayesian modelling showed that ageing did not affect the ability to arbitrate between integration and segregation under either unspeeded or speeded conditions. However, modelling the within-trial dynamics of evidence accumulation under speeded conditions revealed that older observers accumulate noisier auditory representations for longer, set higher decisional thresholds, and have impaired motor speed. Older observers preserve audiovisual localisation performance, despite noisier sensory representations, by sacrificing response speed.

## 1. Introduction

Throughout life we are continually exposed to a barrage of sensory signals. Our ability to effectively navigate through and respond to the world requires us to merge information from multiple sensory modalities into a coherent percept. We may, for example, more easily locate a predator in thick foliage by combining the sight of its movement with the sound of footsteps.

Accumulating evidence suggests that ageing affects how observers integrate sensory signals into perceptual decisions. In speeded target detection paradigms older adults show greater multisensory response facilitation (i.e. redundant target effect; Laurienti et al., 2006; Mahoney et al., 2011). Further, older participants have been shown to integrate multisensory stimuli differently in illusionary settings such as the sound-induced flash illusion (DeLoss et al., 2013; McGovern et al., 2014; Setti et al., 2011) and the McGurk-MacDonald effect (Sekiyama et al., 2014; Setti et al., 2013). Yet, the computational mechanisms underlying these age differences in multisensory integration remain unclear.

Two key mechanisms need to be distinguished: First, ageing is known to reduce the reliability of auditory and visual representations (Dobreva et al., 2011; Lindenberger & Baltes, 1994; Otte et al., 2013; Salthouse et al., 1996). Differences in the reliability of sensory representations may in turn alter the weights that are assigned to the sensory signals during the integration process, thereby changing the final percept. Further, less reliable sensory representations will also reduce observers’ ability to determine whether sensory signals come from a common source and thereby influence how they arbitrate between sensory integration and segregation. In short, age-related increases in noise in the unisensory representations may alter the perceptual outcome of multisensory integration, even if the integration processes are intact.

Second, ageing may genuinely impact how observers arbitrate between sensory integration and segregation depending on temporal, spatial or higher-order statistical correspondence cues or how they weight sensory signals in the integration process. As a consequence, even if unisensory processing were preserved, we would observe differences in multisensory perception.

In short, both changes in unisensory representations and multisensory integration can alter perceptual outcomes in a similar fashion. We thus need to apply models that allow us to dissociate between those two mechanisms.

In the laboratory, the computational principles of multisensory integration have been studied extensively in spatial ventriloquist paradigms where observers need to report their perceived sound (or visual) location when presented with synchronous, yet spatially disparate, auditory and visual signals. For small spatial disparities, observers’ perceived sound location is shifted (or biased) towards the location of the visual signal and vice versa depending on the relative auditory and visual reliabilities—a phenomenon known as the spatial ventriloquist effect. Yet, for large audiovisual spatial disparities where it is unlikely that signals come from a common source, audiovisual interactions and crossmodal biases are attenuated. Recent psychophysics and neuroimaging studies have shown that younger observers arbitrate between sensory integration and segregation in a way that is consistent with the predictions of hierarchical Bayesian Causal Inference (BCI; Aller & Noppeney, 2019; Koerding et al., 2007; Rohe, Ehlis, & Noppeney, 2019; Rohe & Noppeney, 2015a, 2015b; Shams & Beierholm, 2010; Wozny et al., 2010). Bayesian Causal Inference enables arbitration between sensory integration and segregation by explicitly modelling the two causal structures (i.e. common or independent causes) that could have generated the sensory signals. If signals are caused by the same source they are integrated, weighted in proportion to their relative sensory reliabilities; if they are caused by different sources they are treated separately. To account for observers’ uncertainty about the world’s causal structure, a final estimate (e.g. an object’s location) is obtained by averaging the estimates under the assumptions of common and independent sources weighted by their respective posterior probabilities, a decision strategy referred to as model averaging (for other decision functions see Wozny et al., 2010). Spatial ventriloquism, together with Bayesian Causal Inference, may thus allow us to tease apart whether ageing affects only sensory reliabilities (i.e. sensory variance) or also observers’ multisensory binding (as quantified by the model’s causal prior), and to test whether older adults still respond in a way that is consistent with the predictions of BCI.

However, current models of Bayesian Causal Inference do not account for temporal constraints imposed by our natural world and the dynamics of observers’ perceptual inference; BCI enables predictions only for an observer’s response choices (e.g. spatial localisation) but not for his or her response times. In our natural environment we often need to trade off accuracy for speed: a faster, less accurate estimate of the location of a predator may prove far more useful than a highly accurate but slow one. Indeed, recent studies have shown that putatively suboptimal multisensory behaviour can be considered optimal when the dynamics of perceptual decision making, based on both response choices and times, are taken into account (Drugowitsch et al., 2014). Considering response choices and times together is particularly relevant for understanding the impact of ageing on multisensory integration, as older adults have previously been shown to favour accuracy over speed to a greater degree than younger observers (Smith & Brewer, 1995; Starns and Ratcliff, 2010).

Combining psychophysics and computational modelling, the current study was thus designed to investigate how ageing impacts the computational parameters governing multisensory decision making in both unspeeded and speeded contexts (Koerding et al., 2007; Rohe & Noppeney, 2015a, 2015b; Wozny et al., 2010).

First, in an unspeeded spatial ventriloquist paradigm younger and older observers located the source of a sound (which *implicitly* relies on causal inference; see above) or judged whether the auditory and visual signal originated from the same source (which *explicitly* requires the observer to infer the causal structure underlying the audiovisual signals). We assessed how ageing affects observers’ auditory and visual reliabilities (i.e. sensory noise), spatial prior (i.e. spatial expectations), and causal prior (i.e. multisensory binding tendency), as key parameters of the Bayesian Causal Inference model.

Second, in a speeded spatial ventriloquist paradigm observers were presented with spatially congruent or incongruent audiovisual signals and rapidly discriminated whether the auditory (or visual) stimulus was presented in their left or right hemifield. We used a modified version of the Bayesian compatibility bias model (Noppeney, Ostwald, & Werner, 2010; Yu et al., 2009) to characterise how observers accumulate evidence concurrently about signal location and audiovisual spatial congruency (i.e. causal structure), and to make predictions jointly for response choices and times. The age groups were compared in terms of auditory and visual reliabilities, prior binding tendency, and final response threshold.

If older observers differ from younger observers only in sensory reliabilities in unspeeded and speeded contexts, age-related changes in perceptual outcomes are a consequence of their noiser sensory representations. However, if older observers’ behaviour is inconsistent with principles of Bayesian Causal Inference or explained by increases or decreases in their multisensory binding tendencies (as quantified by the causal prior), then ageing genuinely impacts multisensory interactions.

## 2. Methods

### 2.1. Participants

Twenty-three younger adults (eleven male, mean age = 19.5, *SD* = 1.6, range = 18 – 26 years) and twenty-three older adults (seven male, mean age = 72, *SD* = 5.2, range = 63 – 80 years) were included in the study. One older adult was excluded before testing was completed as she was unable to perform unisensory auditory localisation (approximately the same response was given to all auditory stimuli, regardless of source location). The younger adults were undergraduate psychology students at the University of Birmingham, and were compensated in cash or course credits for their time. Older adults were recruited to the study from a database of local participants maintained by the University of Birmingham’s School of Psychology, and were compensated in cash. These community-living older adults had a diverse range of backgrounds; 39% reported education at degree level or above. All participants reported normal hearing and normal or corrected-to-normal vision, and were screened for basic auditory and visual localisation ability using a forced left/right discrimination task (see Supplementary S1). Participants gave informed consent prior to the commencement of testing. The research was approved by the University of Birmingham Ethical Review Committee.

### 2.2. Experimental Setup

Participants were seated at a chin rest 130 cm from a sound-transparent projector screen. Behind the screen, at the vertical centre, a shelf held an array of nine studio monitors (Fostex PM04n) spaced horizontally by 7° of visual angle, including a speaker in the middle of the screen. Auditory stimuli were presented via these speakers at approximately 75 dB SPL. The locations of the speakers were not known to participants. Images were displayed using a BENQ MP782ST multimedia projector at a total resolution of 1280 x 800. All stimuli were presented using The Psychophysics Toolbox 3 (Kleiner, Brainard, & Pelli, 2007) in MATLAB R2010b running on a Windows 7 PC.

Responses were made using a two-button response pad or optical mouse, and in all cases this was effectively self-speeded; the next trial would not begin until a valid response was made. However, for the speeded ventriloquist task it was emphasised to participants that they should respond as quickly as possible while maintaining accuracy. See Figure 1A for an outline of the setup.

**Figure 1.**
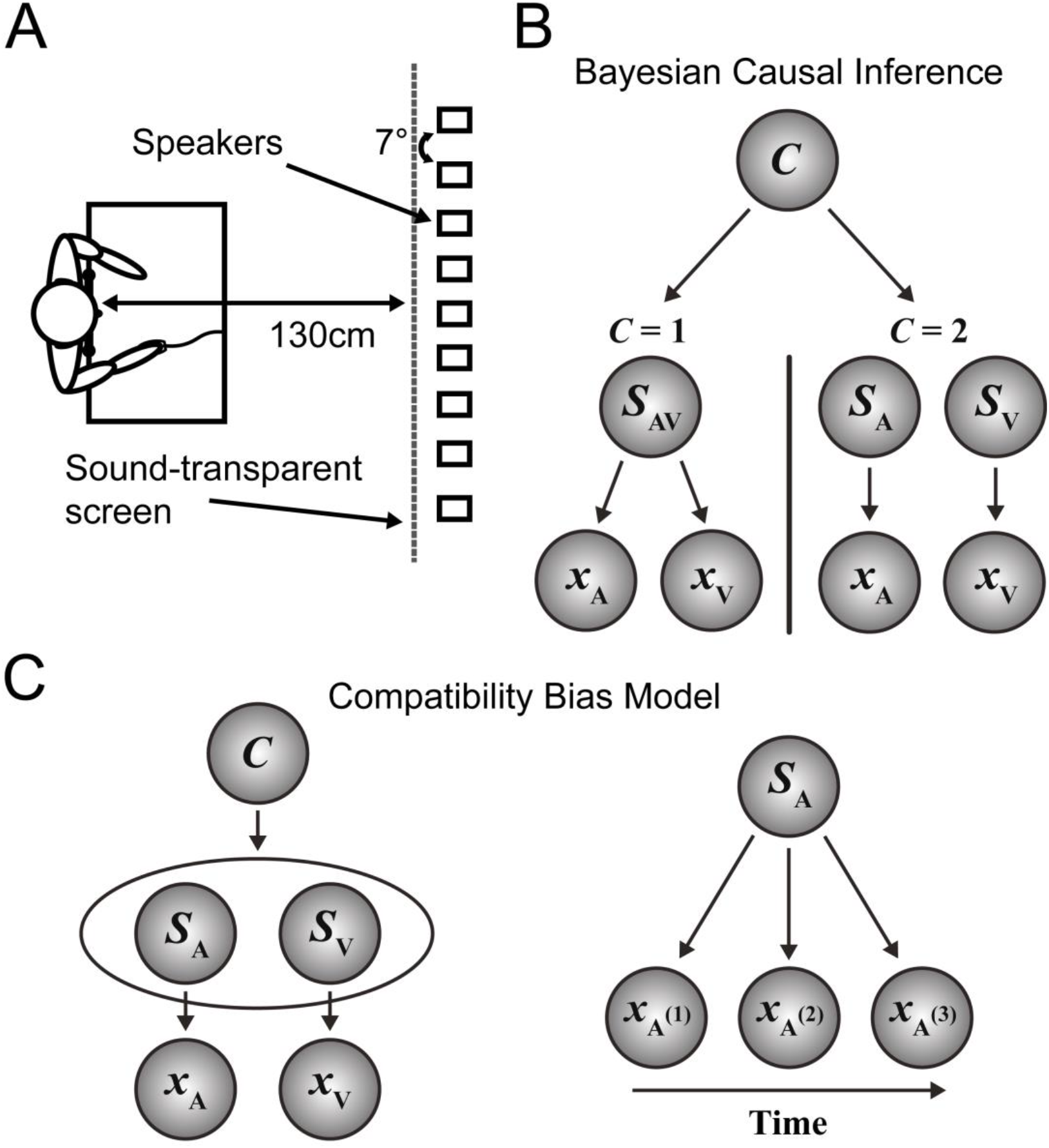
Experimental setup and generative models. (A) Participants were presented with visual stimuli on a sound-transparent projector screen. Sounds were produced by individual speakers concealed behind this screen, which were separated by 7° of visual angle. Responses were given via a mouse or a two-button response pad. (B) Bayesian Causal Inference (BCI) model, based on Koerding et al. (2007). Auditory (*x*_*A*_) and visual (*x*_*V*_) signals may be generated by one common (*C* = 1) audiovisual source (*S*_*AV*_), or by separate (*C* = 2) auditory (*S*_*A*_) and visual (*S*_*V*_) sources. (C) Compatibility bias model, adapted from Yu et al. (2009). Left: Auditory (*S*_*A*_) and visual (*S*_*V*_) sources can either be congruent (*C* = 1, i.e. in same hemifield) or incongruent (*C* = 2, i.e. in opposite hemifields). Right: Across time, the auditory source generates a series of auditory inputs, and the visual source (not shown) a series of visual inputs, in an independent and identical fashion.

### 2.3. Stimuli

Visual stimuli consisted of a 50 ms flash of 15 white (88 cd/m^2^) dots, each 0.44° of visual angle in diameter, against a dark grey (4 cd/m^2^) background. Dot locations were sampled uniquely for each trial from a bivariate Gaussian distribution, with a constant vertical standard deviation of 5.4°. The horizontal standard deviation of this dot cloud was varied to manipulate the reliability of spatial information, with a wider cloud (expressed in degrees of visual angle) resulting in less reliable stimuli (Rohe & Noppeney, 2015). We define the specific horizontal standard deviations used for each paradigm below.

The auditory stimulus was a burst of white noise (duration: 50 ms) played from one speaker in the array in synchrony with the visual stimulus. Sounds were generated individually for each trial and ramped on/off over 5ms. Across all tasks participants fixated a central cross (0.22° radius) that was constantly presented throughout the entire experiment.

### 2.4. Unspeeded audiovisual spatial ventriloquist paradigm

#### 2.4.1. Design and procedure

In a spatial ventriloquist paradigm observers were presented with synchronous auditory and visual stimuli at variable audiovisual spatial disparities and performed implicit or explicit causal inference tasks in separate blocks. First, in an auditory selective attention task, observers reported their perceived sound location. As highlighted in the introduction, spatial localisation implicitly relies on solving the causal inference problem. Second, they explicitly inferred and reported the causal structure (i.e. common vs. independent sources) that could have generated the audiovisual signals in common source judgements.

Irrespective of task context, on each trial auditory and visual stimuli were independently sampled from five possible locations (−14°, −7°, 0, 7, or 14°), and could therefore be spatially congruent or incongruent with varying degrees of disparity (0°, 7°, 14°, 21°, or 28°). Visual stimuli had three levels of reliability (horizontal *SD* of 2°, 6° or 16°) (n.b. a fourth level of visual reliability was excluded from the analysis because the dots were erroneously sampled). The paradigm thus conformed to a 5 (A locations) x 5 (V locations) x 3 (V reliabilities) factorial design.

In the sound localisation task participants reported the perceived sound location as accurately as possible, after a 500 ms post-stimulus delay, by moving a mouse-controlled cursor (white, subtending 9° in height and 0.5° wide) whose movement was constrained to the horizontal plane. The next trial was started one second after observers had indicated their perceived auditory location by clicking the mouse button. Trials were presented randomly in 200-trial blocks. In total, participants completed 600 trials (8 [repetitions] x 5 [A locations] x 5 [V locations] x 3 [V reliabilities)]) of this task.

In the common-source judgement task participants reported whether they perceived the auditory and visual signals to have originated from the same location. 500ms after the presentation of the flash and beep, the words “same” and “different” appeared respectively above and below the fixation cross. Participants indicated with a button press whether the sound and flash were generated by a common source. Participants again completed 600 trials (8 [repetitions] x 5 [A locations] x 5 [V locations] x 3 [V reliabilities)]) of this task, delivered in three blocks of 200 trials.

Unisensory auditory or visual localisation blocks were also included to improve estimation of sensory reliabilities. In unisensory auditory blocks, observers were presented with sounds randomly at one of the five locations and indicated their perceived sound location with the mouse cursor, as above. 80 trials of this task (16 per location) were completed in one block. In unisensory visual blocks, stimuli from the three reliability levels indicated above (horizontal *SD* of 2°, 6° or 16°) were presented randomly in one of the five locations and participants instructed to locate the centre of the dot cloud with the mouse cursor. 120 trials of this task (8 per location, per reliability level) were completed in one block.

#### 2.4.2. Bayesian Causal Inference model

We use Bayesian Causal Inference (BCI; Aller & Noppeney, 2019; Koerding et al., 2007; Rohe, Ehlis, & Noppeney, 2019; Rohe & Noppeney, 2015a, 2015b; Shams & Beierholm, 2010; Wozny et al., 2010) to investigate how younger and older observers arbitrate between sensory integration and segregation. In the following we briefly describe the BCI model; for further details see Koerding et al. (2007).

The BCI generative model assumes that common (*C* = 1) or independent (*C* = 2) sources are determined by sampling from a binomial distribution with the causal prior *P*(*C* = 1) = *p*_common_. For a common source, the “true” location *S*_*AV*_ is drawn from the spatial prior distribution *N*(*μ*_*P*_, *σ*_*P*_). For two independent causes, the “true” auditory (*S*_*A*_) and visual (*S*_*V*_) locations are drawn independently from this spatial prior distribution. For the spatial prior distribution, we assumed a central bias (i.e. *μ*_*P*_ = 0). We introduced sensory noise by drawing *x*_*A*_ and *x*_*V*_ independently from normal distributions centered on the true auditory (respectively visual) locations with parameters *σ*_*A*_ (respectively *σ*_*V*_ for each visual reliability level).

Thus, the generative model included the following free parameters: the causal prior *p*_*common*_, the spatial prior standard deviation *σ*_*P*_, the auditory standard deviation *σ*_*A*_, and visual standard deviations corresponding to the three visual reliability levels *σ*_*V1*_, *σ*_*V2*_, and *σ*_*V3*_.

During perceptual inference the observer is assumed to invert this generative model. The probability of the underlying causal structure can be inferred by combining the causal prior with the sensory evidence according to Bayes’ rule:

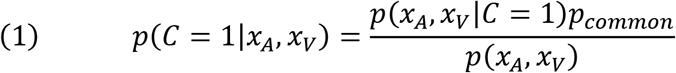

We assumed that subjects report ‘common source’ (i.e. explicit causal inference) when the posterior probability of a common source is greater than the threshold of 0.5:

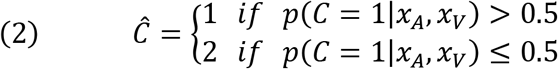

In the case of a common source (*C* = 1; Figure 1B left), the maximum a posteriori probability estimate of the auditory location is a reliability-weighted average of the auditory and visual estimates and the prior.

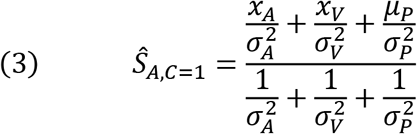

In the case of a separate-source inference (*C* = 2; Figure 1B right), the estimate of the auditory signal location is independent from the visual spatial signal.

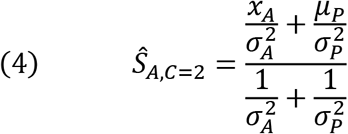

Given the decisional strategy of model averaging (for other decisional strategies see Wozny et al., 2010) the observer will compute a final auditory localisation estimate by averaging the spatial estimates under common and independent source assumptions, weighted in proportion to their posterior probabilities (i.e. implicit causal inference).

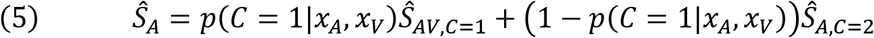

The predicted distributions of the auditory spatial estimates, *p*(*Ŝ*_*A*_|*S*_*A*_, *S*_*V*_), and the common source estimates, *p*(*Ĉ*|*S*_*A*_, *S*_*V*_), were obtained by marginalising over the internal variables *x*_*A*_ and *x*_*V*_. For the unisensory auditory and visual localisation tasks, we used the predicted distributions *p*(*Ŝ*_*A,C*=2_|*S*_*A*_) for auditory blocks and *p*(*Ŝ*_*V,C*=2_|*S*_*V*_) respectively.

These distributions were generated by simulating *x*_*A*_ and *x*_*V*_ 10000 times for each of the conditions and inferring *Ŝ*_*A*_, *Ŝ*_*A,C*=2_, *Ŝ*_*V,C*=2_, and *Ĉ* from the equations above. Based on these predicted distributions (given an additional noise kernel with a fixed *σ*_*motor*_ = 1), we computed the log-likelihood of participants’ auditory localisation and common-source judgement responses.

We fitted the Bayesian Causal Inference model jointly to observers’ localisation responses in the audiovisual and the unisensory visual and auditory stimulation conditions. We modelled the sensory noise and spatial prior parameters separately for unisensory and bisensory trials, as this was found to fit the data best overall (see Supplementary S5 for a formal comparison with models that did not separate parameters based on unisensory or audiovisual context). Therefore, a total of eleven free parameters was fitted for each participant: the causal prior *p*_*common*_, the spatial prior standard deviations *σ*_*P uni*_ and *σ*_*P bi*_, the auditory standard deviations *σ*_*A uni*_ and *σ*_*A bi*_, and visual standard deviations corresponding to the three visual reliability levels *σ*_*V1 uni*_, *σ*_*V2 uni*_, *σ*_*V3 uni*_, *σ*_*V1 bi*_, *σ*_*V2 bi*_, *σ*_*V3 bi*_ (indices *uni* and *bi* correspond to unisensory and bisensory trials respectively). Assuming independence of conditions and responses, we summed the log-likelihoods across conditions and across localisation and common-source judgement responses to obtain a single log-likelihood for each subject. To obtain maximum likelihood estimates for each subject’s model parameters we used a Bayesian adaptive search algorithm (BADS; Acerbi & Ma, 2017) with the parameters for initialisation determined by a prior grid search.

The parameters (causal prior, spatial prior[s], and sensory variances) obtained from the winning model were compared between age groups using separate non-parametric Mann-Whitney *U* tests. We also calculated Bayes factors using the Bayesian Mann-Whitney test as implemented in JASP (JASP Team, 2018; van Doorn et al., 2018) using the default Cauchy prior (scale = 0.707).

### 2.5. Speeded ventriloquist paradigm

#### 2.5.1. Design and procedure

To assess participants’ audiovisual integration of spatial cues under speeded conditions, taking into account both final responses and reaction times, we used a simpler 2 (auditory location: left vs. right) x 2 (visual location: left vs. right) x 2 (relevant and reported sensory modality: auditory vs. visual) ventriloquist paradigm. On each trial, a visual stimulus with horizontal *SD =* 5.4° was displayed simultaneously with a burst of white noise. The centre of the visual cloud and the white noise were presented at 14° either left or right of a central fixation cross. These audiovisual stimuli were spatially congruent on half of the trials, and incongruent on the other half. In an auditory or visual selective attention paradigm, participants indicated either the location of the sound (respond-auditory task) or the cloud (respond-visual task) as quickly and accurately as possible via a two-choice key press, while ignoring the other modality. The task was self-speeded in this way (i.e. no response deadline) as any imposed incentives or timing criteria may have affected the groups differently; we rely on the compatibility bias model (Yu et al., 2009; described below) to separate age differences in motor speed and speed/accuracy trade-off from potential differences in sensory reliability/evidence accumulation. The tasks were performed in two blocks of 160 trials. The order of these tasks was counterbalanced between participants. In total the experiment included 320 trials: 40 (repetitions) x 2 (visual location) x 2 (auditory location) x 2 (reported sensory modality).

#### 2.5.2. Compatibility bias model

To assess age differences in responses to multisensory stimuli under temporal constraints, we analysed the respond-auditory data by adapting the “compatibility bias” model to an audiovisual context (Noppeney, Ostwald, & Werner, 2010; Yu et al., 2009). This models the within-trial dynamics of audiovisual evidence accumulation, leading to predictions for both response choice and response times.

See Yu et al. (2009) for full details about the compatibility bias model. Briefly, this generative model assumes that congruent (*C* = 1) or incongruent (*C* = 2) sources are determined by sampling from a binomial distribution with the compatibility or congruency prior *P*(*C* = 1) = *p*_*congruency*_. The visual *S*_*V*_ and auditory *S*_*A*_ sources can either be left (–1) or right (+1). For a congruent trial, the auditory and visual locations are identical, i.e. *S*_*A*_ = *S*_*V*_ (*S*_*A*_ and *S*_*V*_ are either both left or both right). For an incongruent trial, the auditory and visual locations are in opposite hemifields, i.e. *S*_*A*_ = –*S*_*V*_ (two possibilities: *S*_*A*_ = –1 and *S*_*V*_ = 1, or *S*_*A*_ = 1 and *S*_*V*_ = –1). Hence we obtain a total of four possible stimulus combinations. We then sample noisy sensory inputs successively for each time point within a trial by drawing ***x***_*t*_ = [*x*_*A*_(*t*) *x*_*V*_(*t*)] independently from normal distributions centred on *S*_*A*_ (or *S*_*V*_) with parameters *σ*_*A*_ (or *σ*_*V*_ respectively). This thereby models that the brain receives progressively more information about the location of the auditory and visual sources and thus, indirectly, about whether or not they are congruent (n.b. though in our experiment auditory and visual inputs are brief, we model evidence accumulation via feedback loops as a series of sensory inputs). Based on a stream of audiovisual inputs ***X***_*t*_ = [***x***_1_, ***x***_*2*_, ***x***_*3*_ … ***x***_*t*_] the observer is then assumed to compute the posterior probability over congruency *C* and auditory (or visual) source location iteratively according to Bayes’ rule (initialised with the prior *P*(*C*) = *β*):

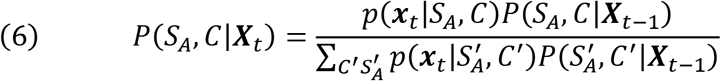

A left/right decision is then made when the evolving trajectory of the marginal

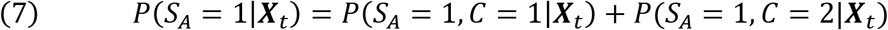

reaches a threshold *q*.

Thus, incongruent visual information should be most influential on perceived auditory location at the onset of the trial, when the initial compatibility prior dominates, but this influence decreases as information about the location of each stimulus is accumulated. The process is terminated when sufficient evidence is accumulated about the location of the auditory stimulus for a decisional threshold to be reached, after which a left/right spatial response is made. To accommodate that older adults have slower motor speed than younger adults (as confirmed by a separate finger tapping task reported in Supplementary S2), we included an additional non-decision-time parameter *t*_nd_ to account for motor delays.

The model therefore has five free parameters in total: the compatibility prior (i.e. prior probability of audiovisual signals coming from a common cause) *β*; the standard deviations of the auditory and visual signals, *σ*_*A*_ and *σ*_*V*_ respectively; the response threshold *q*; and a non-decision-time parameter *t*_*nd*_ that allows for a variable motor delay between the threshold being reached and a response being given.

As in the Bayesian Causal Inference model we obtained the predicted distributions of the auditory spatial estimates, *P*(*Ŝ*_*A*_|*S*_*A*_, *S*_*V*_), and response times, 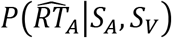, by marginalising over the internal variables *x*_*A*_ and *x*_*V*_. These distributions were generated by simulating *x*_*A*_ and *x*_*V*_ for 300 time steps (of 10 ms length) 10000 times for each of the conditions. For each simulated trial with a series of 300 *x*_*A*_ and *x*_*V*_, we then computed the response time and choice when *P*(*S*_*A*_ = −1|***X***_*t*_) first crossed the decisional threshold *q* using Equations 5 and 6 above. Based on these predicted response choice and response time distributions, we computed the log-likelihood of participants’ auditory (or visual) localisation responses and the response times (after adding the non-decision time *t*_*nd*_). Assuming independence of conditions as well as independence of the log-likelihoods for response times and choices, we summed the log-likelihoods across conditions and across response times and choices for a particular subject. To obtain maximum likelihood estimates for the model parameters for each subject (*β*, *σ*_*A*_, *σ*_*V*_, *q*, *t*_*nd*_) we used a Bayesian Adaptive Search optimisation algorithm (BADS; Acerbi & Ma, 2017) with parameters initialised based on a grid search.

To investigate whether any of the parameters of these two Bayesian models were significantly different between older and younger adults the fitted parameters were entered into separate non-parametric Mann-Whitney *U* tests. We also calculated Bayes factors using the Bayesian Mann-Whitney test as implemented in JASP (JASP Team, 2018; van Doorn et al., 2018) using the default Cauchy prior (scale = 0.707).

## 3. Results

### 3.1. Unisensory screening tests and the Montreal Cognitive Assessment

Prior to the main unspeeded and speeded ventriloquist experiments, all observers were screened for basic auditory and visual localisation ability with a binary left/right forced-choice spatial classification task. Individuals were characterised in terms of the slope and threshold of psychometric functions fitted to these responses. Older and younger adults were closely matched: no significant age differences in threshold or bias were observed for auditory or visual spatial processing, suggesting that sensory spatial reliability was approximately similar between age groups. No participants were excluded as a result of poor performance on this task. See Supplementary S1 for full details.

Older participants were also screened using the Montreal Cognitive Assessment with a cut-off score of 23 (Coen et al., 2011; Roalf et al., 2013; Luis et al., 2009); none of our older participants scored below 25.

### 3.2. Unspeeded ventriloquist paradigm: Localisation and common source judgement

#### 3.2.1. Descriptive and GLM-based analysis

An unspeeded spatial ventriloquist paradigm was used to compare younger and older adults’ responses to audiovisual spatial stimuli in the absence of temporal constraints. Figure 2 shows participants’ auditory localisation (presented in terms of the magnitude of ventriloquist effect, *VE* = [*A*_*resp*_ – *A*_*loc*_] / [*V*_*loc*_ – *A*_*loc*_]) and common-source judgement responses (characterised as the probability of responding “same-source”) as a function of visual reliability level and audiovisual disparity. As predicted by Bayesian Causal Inference, the ventriloquist effect was strongest when visual reliability was high and the audiovisual disparity small. The age groups performed remarkably similarly on both measures, with standard GLM analyses revealing no significant effects of age on final response choices. However, older observers were significantly slower than younger adults when localising sounds in the spatial ventriloquist paradigm. Further, we observed significant age effects on the common-source judgement reaction times (Figure 2D), including significant interactions between age, visual reliability, and audiovisual disparity. See Supplementary S3 for full GLM analyses of these results.

**Figure 2.**
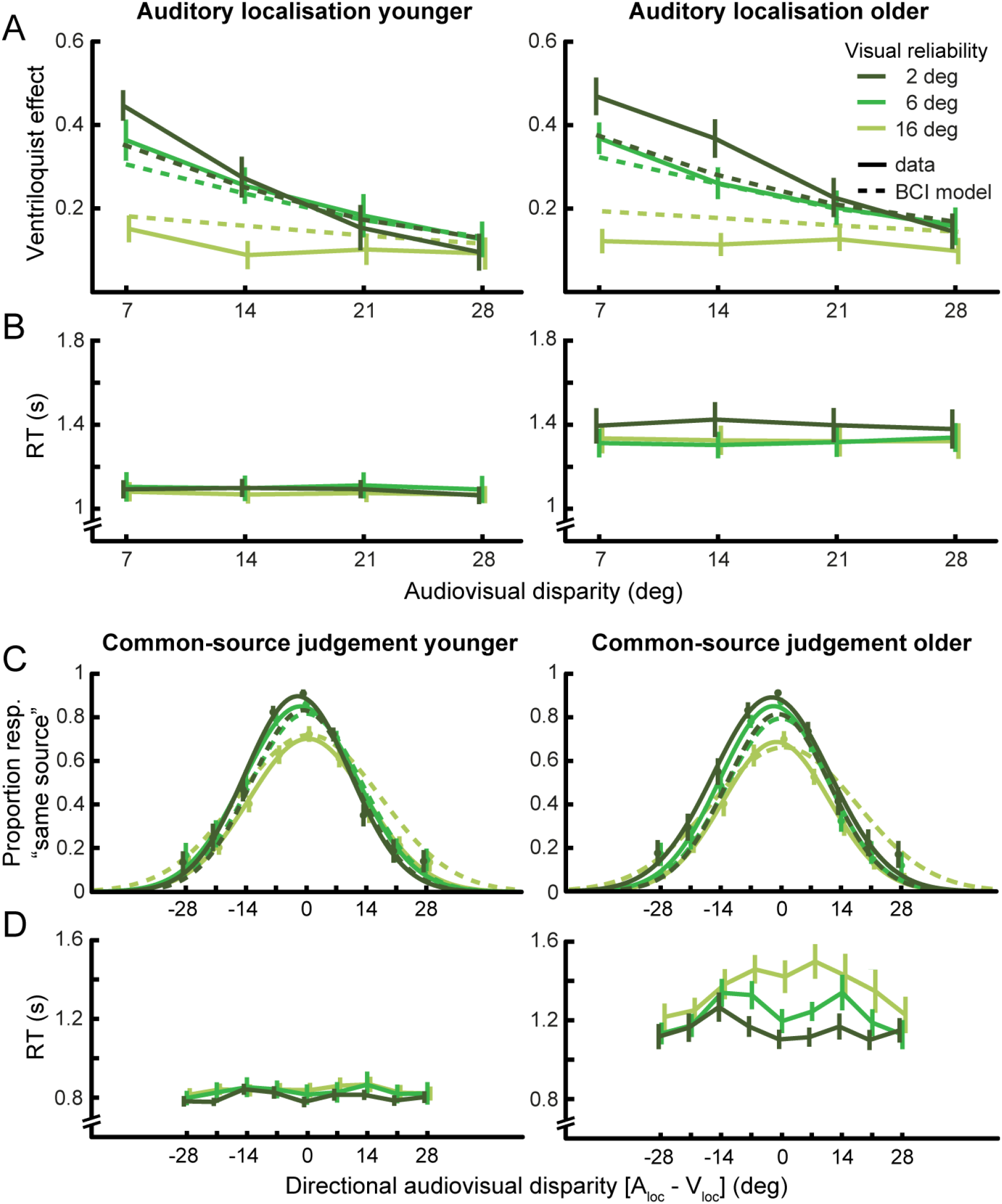
Behavioural responses, reaction times and BCI model predictions for younger and older adults. (A) Relative ventriloquist effect (*VE* = [*A*_*resp*_ – *A*_*loc*_] / [*V*_*loc*_ – *A*_*loc*_]) for auditory localisation, shown as a function of audiovisual disparity (*x*-axis, pooled over direction) and visual reliability (colour coded). Behavioural data (mean across subjects, solid lines) and the predictions of the Bayesian Causal Inference model (dashed lines) are shown. (B) Reaction times in auditory localisation task. (C) Proportion reported “same source” in common-source judgement task, as a function of audiovisual disparity and visual reliability. The panels show the Gaussians fitted to the behavioural response (mean across subjects, solid lines) and the predictions of the Bayesian Causal Inference model (dashed lines). (D) Reaction times (pooled over response; mean across subjects) in common-source judgement task. Error bars show ±1 *SEM*.

#### 3.2.2. Bayesian modelling

Table 1 summarises the fitted parameters (within-group mean and *SD*) of the Bayesian Causal Inference model for younger and older participants. Table 1 also reports the results of the nonparametric tests comparing the parameters between the older and younger groups together with the Bayes factors associated with each statistical comparison. We observed small but significant group differences in auditory and visual variance parameters that were estimated based on unisensory localisation tasks alone, suggesting that older adults were slightly less precise when locating both auditory and particularly unreliable visual stimuli. These group differences were not significant when the sensory variance parameters were estimated based on responses to audiovisual stimuli, probably because these parameters were less precisely estimated in this case: in the audiovisual context the visual variance parameter is only estimated indirectly from auditory responses, and the auditory variance is always estimated in the presence of interfering visual signals (and so may be influenced by factors other than peripheral sensory noise).

**Table 1.** Bayesian Causal Inference parameters (across-participants mean, *SD*) for younger (*n* = 23) and older (*n* = 22) participants. Mann-Whitney *U* tests with Bayes factors comparing the BCI parameters between older and younger adults. The Bayesian Causal Inference model was fitted jointly to unisensory and audiovisual conditions allowing for separate parameters for the standard deviation of the spatial prior (*σ*_*P,uni*_, *σ*_*P,bi*_) and sensory noise (*σ*_*A,uni*_, *σ*_*A,bi*_, *σ*_*V1,uni*_, *σ*_*V1,bi*_, … *σ*_*V3,uni*_, *σ*_*V3,bi*_). The standard deviation of motor response *σ_motor_* was constrained to be the same for unisensory and multisensory localisation responses. *BF*_10_ quantifies degree of support for the alternative hypothesis that the groups differ, relative to the null hypothesis; *BF*_01_ shows degree of support for the null hypothesis that there is no difference between groups, relative to the alternative hypothesis.

| | Younger | | Older | | Mann-Whitney $U$ | | | Bayes factors | |
| --- | --- | --- | --- | --- | --- | --- | --- | --- | --- |
| | Mean | $SD$ | Mean | $SD$ | $W$ | $p$ | $\eta^2$ | $BF_{10}$ | $BF_{01}$ |
| Unisensory |  |  |  |  |  |  |  |  |  |
| $\sigma_{P\ uni}$ | 37.20 | 35.69 | 24.79 | 28.63 | 299 | .305 | .02 | 0.46 | 2.16 |
| $\sigma_{A\ uni}$ | 5.27 | 1.96 | 6.79 | 2.76 | 155 | **.026** | .11 | 2.19 | 0.46 |
| $\sigma_{V1\ uni}$ | 1.76 | 1.22 | 2.10 | 1.06 | 174 | .075 | .07 | 1.10 | 0.91 |
| $\sigma_{V2\ uni}$ | 2.32 | 0.76 | 2.89 | 1.52 | 198 | .218 | .04 | 0.54 | 1.84 |
| $\sigma_{V3\ uni}$ | 4.22 | 1.00 | 5.38 | 1.67 | 132 | **.005** | .17 | 4.95 | 0.20 |
| Bisensory |  |  |  |  |  |  |  |  |  |
| $P_{common}$ | 0.42 | 0.13 | 0.43 | 0.13 | 245 | .866 | < .01 | 0.30 | 3.28 |
| $\sigma_{P\ bi}$ | 38.71 | 25.88 | 32.20 | 27.37 | 303 | .264 | .03 | 0.40 | 2.49 |
| $\sigma_{A\ bi}$ | 8.59 | 4.40 | 9.37 | 5.78 | 234 | .677 | < .01 | 0.35 | 2.84 |
| $\sigma_{V1\ bi}$ | 3.19 | 4.08 | 3.08 | 3.13 | 241 | .796 | < .01 | 0.30 | 3.31 |
| $\sigma_{V2\ bi}$ | 5.12 | 4.32 | 6.07 | 5.47 | 204 | .274 | .03 | 0.44 | 2.25 |
| $\sigma_{V3\ bi}$ | 12.79 | 9.72 | 20.61 | 26.13 | 209 | .327 | .02 | 0.48 | 2.09 |

Crucially, however, no significant group differences were observed for the *P*_*common*_ or *σ*_*P*_ parameters. This suggests that the two age groups had similar central spatial priors and causal priors, suggesting that older and younger adults showed similar tendencies to bind audiovisual signals (in an unspeeded context) consistent with Bayesian Causal Inference.

To verify that these results were not confounded by possible age differences in motor noise (i.e. noisier mouse localisation responses), we also fitted a version of the model that allowed the parameter *σ*_*motor*_ to vary freely (*σ*_*motor*_ was fixed at 1° for all participants in the main analysis). The pattern of results remained similar, though the group difference in *σ*_*A uni*_ became marginally non-significant (*p* = .052). Further, there were no significant group differences in the *σ*_*motor*_ parameter (*p* > .05, *BF*_*01*_ = 3.15). See Supplementary S6 for details.

In summary, age did not influence observer’s implicit (auditory localisation) or explicit (common-source judgement) causal inference in terms of response choices. Our Bayesian modelling analysis revealed that older adults had slightly noisier auditory and visual representations when estimated separately for the unisensory conditions. Importantly, though, the comparable causal prior (and central prior), and similar mean response choices, indicate that older observers combined audiovisual spatial signals according to the same computational principles as younger adults.

Yet, ageing was associated with complex changes in reaction times to multisensory stimuli. The profile of these age differences suggests that older adults took more time to respond when the causal structure of the stimuli was more ambiguous and the task therefore more challenging, such as when the visual stimulus was less reliable and/or the audiovisual disparity of intermediate size. These response time findings were followed up in a speeded ventriloquist task, where observers were explicitly instructed to respond as quickly as possible while maintaining accuracy.

### 3.3. Speeded ventriloquist paradigm

#### 3.3.1. Descriptive and GLM-based analysis

A simplified, speeded ventriloquist paradigm was used to assess younger and older adults’ responses to audiovisual spatial stimuli under speed instructions. Figure 3 summarises response accuracy (panel B) and speed (panel C) for younger and older adults; trials are pooled over left and right to characterise them in terms of spatial (in)congruence. Standard GLM analysis of these results shows that older adults were significantly more accurate than younger adults in the respond-visual task. Older adults were also significantly slower overall and, importantly, age interacted with congruence in the respond-auditory tasks (see Section 3.2). Mirroring the profile of the unspeeded common-source judgement responses, older adults again took disproportionately longer to respond under the most challenging conditions where they located the auditory signal in the presence of an incongruent visual distractor. See Supplementary S4 for full GLM analysis.

**Figure 3.**
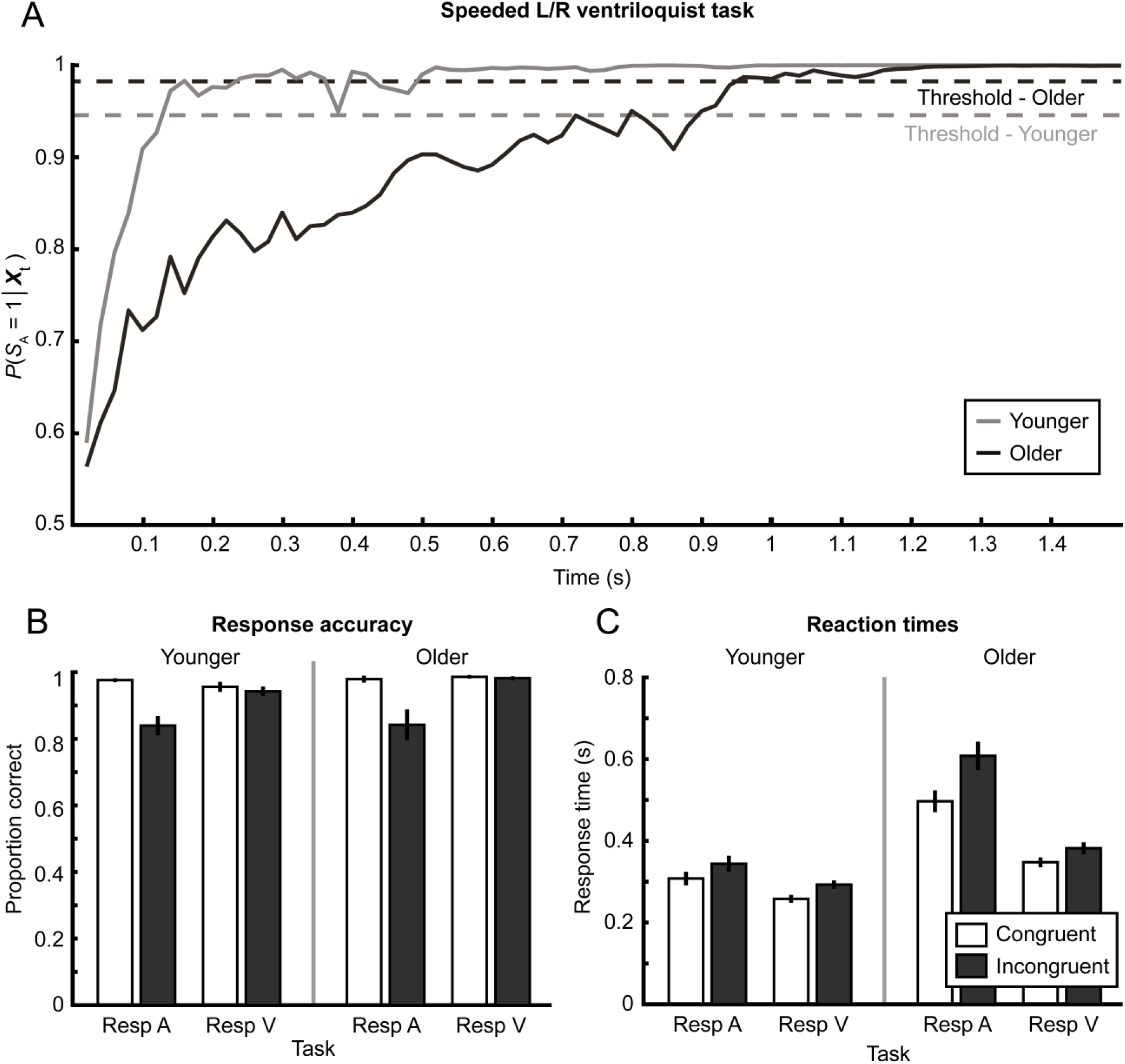
Speeded left/right ventriloquist paradigm and compatibility bias model. (A) Accumulation of evidence traces for the compatibility bias model: for ‘respond auditory’ trials the observer is thought to accumulate audiovisual evidence about whether the auditory source is left = −1 or right = 1 within a trial until a decisional threshold is reached and a response elicited. Solid lines show the posterior probability *P*(*S*_*A*_ = 1|***X***_*t*_) as a function of within-trial time with auditory and visual inputs arriving every 10 ms. Each trace represents the mean across ten (incongruent, auditory right) simulated trials for a representative participant in each group, using these participant’s maximum likelihood parameters. Dashed lines indicate the participants’ fitted decisional thresholds. Older observers accumulate noisier evidence until a higher decisional threshold is reached. (B and C) Response accuracy and reaction times (across-participants mean ± 1 *SEM*) for respond-auditory and respond-visual tasks, separated by spatial congruence (i.e. pooled over left and right).

#### 3.3.2. Compatibility bias model

The compatibility bias model was fitted to participants’ auditory spatial responses and reaction times. This allowed us to characterise how younger and older observers accumulate audiovisual evidence about spatial location and audiovisual congruency until a decisional threshold is reached and a response given. Fitted parameters were compared using separate Mann-Whitney *U* tests and the Bayesian version of the Mann-Whitney test (JASP Team, 2018; van Doorn et al., 2018). See Table 2 for a summary of results.

**Table 2.** Compatibility bias parameters (across-participants mean, *SD*) for younger (*n* = 23) and older (*n* = 22) participants. Mann-Whitney *U* tests with Bayes factors comparing the compatibility bias parameters between older and younger adults: standard deviation of the auditory signal *σ*_A_, standard deviation of the visual signal *σ*_V_, compatibility prior *β*, response threshold *q*, and non-decision-time *t*_nd_. *BF*_10_ quantifies degree of support for the alternative hypothesis that the groups differ, relative to the null hypothesis; *BF*_01_ shows degree of support for the null hypothesis that there is no difference between groups, relative to the alternative hypothesis.

| | Younger | | Older | | Mann-Whitney $U$ | | | Bayes factors | |
| --- | --- | --- | --- | --- | --- | --- | --- | --- | --- |
| | Mean | $SD$ | Mean | $SD$ | $W$ | $p$ | $\eta^2$ | $BF_{10}$ | $BF_{01}$ |
| $\sigma_A$ | 1.53 | 0.54 | 2.93 | 4.01 | 164 | **.044** | .09 | 3.11 | 0.32 |
| $\sigma_V$ | 1.85 | 3.50 | 0.87 | 0.97 | 283 | .507 | .01 | 0.44 | 2.27 |
| $\beta$ | 0.75 | 0.12 | 0.78 | 0.13 | 192 | .169 | .04 | 0.56 | 1.79 |
| $q$ | 0.93 | 0.05 | 0.95 | 0.07 | 141 | **.010** | .14 | 2.68 | 0.37 |
| $t_{nd}$ | 0.22 | 0.05 | 0.33 | 0.07 | 54 | **< .001** | .45 | 5101.52 | < 0.01 |

Corroborating the findings of the BCI model, the age groups did not differ in their prior tendency to integrate multisensory stimuli, quantified in this case by the compatibility prior *β*. However, similar to the results from unspeeded localisation, the auditory signal (*σ*_auditory_) was significantly noisier in older than younger adults, leading to a slower accumulation of evidence and thus (in combination with the motor slowing and higher decision threshold, see below) slower response times. This indicates that it takes older participants longer than their younger counterparts to reach any given level of evidence about the location of an auditory stimulus. The groups did not differ in the variance of the visual input *σ*_visual_. However, the remaining two parameters were also significantly different between the groups. First the non-decision time *t*_nd_, which captures the time between a decision is made and the response given, was significantly higher for the older age group. This is unsurprising; our older adults’ impaired motor speed is confirmed by a separate finger-tapping task reported in Supplementary S2. Second, older adults also set their decision threshold *q* significantly higher, requiring more evidence before deciding on a response. See Figure 3A for an illustration of the model. Taken as a whole, our Bayesian modelling analysis confirms that older adults show a similar multisensory binding tendency and combine signals to the same computational principles as younger adults. However, older adults have noisier unisensory auditory spatial representations. As a result of i. those noisier auditory spatial representations, ii. a different speed-accuracy trade off (i.e. decision threshold *q*) and iii. slower motor speed (i.e. non-decision time *t*_nd_) they have slower response times.

## 4. Discussion

This study investigated the effects of ageing on audiovisual integration for spatial localisation under both speeded and unspeeded conditions. Our results demonstrate that ageing does not fundamentally impact how observers integrate auditory and visual spatial signals into representations of space: older adults showed the same audiovisual binding tendency as the younger age group, and their behaviour conformed similarly to the predictions of the Bayesian models. However, older adults showed noisier sensory, in particular auditory, representations. Moreover, they used a higher decisional threshold, trading off speed for accuracy. This suggests that older observers preserve audiovisual localisation performance, despite noisier sensory representations, by sacrificing response speed.

These results may initially seem surprising in light of accumulating research showing that ageing alters multisensory integration. For example, older adults have been shown to be more susceptible to the sound-induced flash illusion (DeLoss et al., 2013; McGovern et al., 2014; Setti et al., 2011) and to respond differently to McGurk-MacDonald stimuli (Sekiyama et al., 2014; Setti et al., 2013). It is possible, however, for such effects to occur in the absence of age differences in the actual computational processes underlying multisensory perception. Any change that leads to an increase in sensory variances may make the arbitration between common and separate sources more challenging, and/or change the relative weighting of the sensory modalities in the final percept. Potentially, susceptibility to the sound-induced flash illusion is changed with age because it relies on precise representations of stimulus timing that have been shown to be impaired by ageing (Chan et al., 2014; Mazelová et al., 2003). Ng and Recanzone (2017) provide a possible mechanism for this decline: a study of neural responses to simple stimuli in macaque primary auditory cortex found that aged monkeys showed firing patterns that were noisier (i.e. less temporally precise) and less selective than those seen in younger animals. Age-related differences in perception of McGurk-MacDonald stimuli may also be due in part to impaired temporal perception, as the fine temporal structure of speech signals is an important cue for comprehension (especially in the context of competing noise; Moore, 2008). In this case the effect is likely to be further compounded by reductions in speech comprehension, resulting from presbycusis that particularly affects higher sound frequencies (Pichora-Fuller & Souza, 2003). These mechanisms are notably unisensory, and do not imply any change in the computational process of multisensory integration itself.

The argument that older adults’ changed multisensory perception results primarily from differences in unisensory variances, and not from alterations in the computational mechanisms *per se*, can also explain why we did not find significant age differences in the final responses to our multisensory tasks: our unisensory results, and those of others (Dobreva et al., 2011; Otte et al., 2013), demonstrate only limited age differences in localisation ability. Based on screening tests involving binary left/right judgements, younger and older adults were similar in their ability to locate unisensory auditory and visual stimuli. The sensory variance parameters of a Bayesian Causal Inference model fitted to multisensory localisation and common-source judgement responses also did not differ between age groups. However, the same parameters fitted using the more sensitive unisensory free-localisation responses did reveal small but significant age differences in sensory variances, suggesting that older adults were less reliable in their localisation of both auditory and (low-reliability) visual stimuli.

Existing literature is similarly ambiguous about age-related declines in (especially) auditory localisation. Dobreva et al. (2011) report limited but significant age differences in observers’ ability to freely localise transient broadband stimuli along the azimuth, while Otte et al. (2013) found no such effects. It therefore seems that the effects of normal, healthy ageing on auditory localisation ability may be subtle and difficult to detect.

In terms of visual localisation, we note that our older adults are likely to have had impaired accommodation responses compared to the younger age group (Glasser & Campbell, 1997). Depending on the corrective lenses worn (participants were instructed to wear their normal spectacles for testing), this may have led to the older group expending more effort to keep the visual stimuli in focus and/or the stimuli appearing less focused. The small but significant age differences we observed in unisensory visual localisation may be, in part, a reflection of this reduced accommodation ability.

In light of these limited age differences in audiovisual localisation performance, it would be interesting for future research to apply computational modelling to multisensory contexts where strong age differences have been shown previously. The sound-induced flash illusion is a strong candidate for this, as older adults are known to be significantly more susceptible (DeLoss et al., 2013; McGovern et al., 2014; Setti et al., 2011) and young observers’ perception of the illusion has previously been successfully modelled using a BCI framework (Shams et al., 2005). Fitting the BCI model to younger and older observers’ responses would allow us to distinguish whether age differences in perception of the sound-induced flash illusion result from changes in unisensory variances (i.e. noise) or in observers’ multisensory binding itself.

Our discussion of age differences in multisensory integration has thus far addressed only final response choices, ignoring reaction times, but our natural environment does not afford us infinite time to react to multisensory stimuli. When we define and evaluate multisensory integration performance, it is therefore also important to consider the time taken to respond. In fact, GLM-based analyses of common-source judgement reaction times suggested that older adults took disproportionately longer to respond to audiovisual signals at intermediate levels of spatial disparity, where the underlying causal structure (i.e. common vs. independent sources) was less certain. Such findings imply the presence of differences in the groups’ evidence accumulation and decision-making process, and/or in their speed/accuracy criteria, even in an unspeeded context.

We thus applied a simplified, speeded ventriloquist paradigm to directly address the question of age differences in response times to multisensory spatial stimuli. GLM analyses again showed that older adults were disproportionately slower in the most challenging condition, in this case locating a sound in the presence of an incongruent visual distractor. To characterise the computational processes underlying these differences, it is necessary to move beyond the static BCI model to a dynamical approach that can make predictions jointly about observers’ spatial choices and response times. We thus applied the compatibility bias model (Noppeney, Ostwald, & Werner, 2010; Yu et al., 2009) to participants’ auditory judgement responses in this paradigm.

This model assumes that the observer accumulates auditory and visual evidence about the location of the reported stimulus, and about the causal structure of the signals, until a decisional threshold is reached and a response given. It thereby provides an important perspective on the dynamics of decision making within a trial. Again in this case, the fundamental computations were not affected by healthy ageing. Likewise, older adults’ prior binding tendency was not significantly different from the younger group. However, the compatibility bias model also revealed that older adults responded more slowly than younger adults for three reasons. First, older adults have impaired motor speed, as indexed by the non-decision time variable (and confirmed by a supplementary finger-tapping task; see Supplementary S2). Second, they use a higher response threshold, requiring a greater degree of certainty before a response is given. This is consistent with previous studies of age differences in speed/accuracy trade-off (Smith & Brewer, 1995; Starns and Ratcliff, 2010). Third, the compatibility bias model analysis suggests that the auditory representations are less reliable (i.e. greater auditory variance) in older participants, such that evidence accumulates more slowly (see Figure 3). In other words, the initial auditory representation may be noisier and less reliable for older adults, but older observers can achieve equal performance levels (in terms of final response choices) to younger participants by accumulating this noisy evidence for longer via internal feedback loops.

It is important to note that the Bayesian causal inference model, and other approaches that consider only the observer’s final response, may be less sensitive to these age-related changes in internal sensory noise (though the unisensory localisation data do provide some evidence of small reliability differences). This illustrates how dynamical models that accommodate both reaction times and final response choices can provide critical new insights into evidence accumulation and perceptual decision making.

In conclusion, our results demonstrate that multisensory causal inference is preserved in older adults. However, older observers only maintain this performance by accumulating noisier auditory information over a longer period of time. When combined with well-established changes in motor speed and speed/accuracy trade-off, this leads to significant and nonlinear age differences in reaction times to complex multisensory stimuli during spatial localisation.

## Acknowledgements

This research was funded by the European Research Council (ERC-2012-StG_20111109 multsens) and the MRC-ARUK Centre for Musculoskeletal Ageing Research (CMAR)

## Supplementary Methods and Results

### S1. Unisensory L/R discrimination

We administered simple left/right forced-choice tasks to compare age groups on basic unisensory spatial discrimination ability, and to ensure all participants were able to locate auditory and visual stimuli in space sufficiently well for inclusion in the study. We chose these measures as more directly relevant than, for example, pure tone hearing thresholds (older participants are likely to have some impairment at higher frequencies, but this may not result in any substantial decrease in auditory localisation ability).

Participants’ spatial hearing performance (i.e. bias and reliability/variance of auditory spatial representations) was measured using a forced left/right spatial discrimination task. Individual bursts of white noise were emitted from one of seven locations (−21°, −14°, −7°, 0°, 7°, 14°, or 21°) in a random order. Participants indicated as accurately as possible via key press whether the sound originated from the left or right half of the screen. This task involved one block of 210 trials (30 per location).

Visual spatial perception (i.e. bias and variance/reliability of visual spatial representations) was measured using a similar left/right spatial discrimination task. Visual stimuli with horizontal *SD* = 2° or 25° were randomly presented centred at one of seven locations (−21°, −14°, −7°, 0°, 7°, 14°, or 21°); participants indicated whether the centre of the dot cloud originated from the left or right side of the screen. This task involved two blocks of 210 trials each (total 30 trials per condition).

For each participant and stimulus type, we used the Palamedes toolbox for MATLAB (Prins & Kingdom, 2009) to calculate the slope *α* and threshold *β* of cumulative Gaussians fitted to the proportion of “perceived right” responses as a function of true stimulus location. These parameters were allowed to vary freely, while the lapse parameters *γ* and ***λ*** were constrained to be the same and to fall between 0 and 0.05 (Wichmann & Hill, 2001).

Older and younger adults were closely matched on these tasks. For auditory spatial discrimination, an independent-samples *t*-test revealed no significant effect of age on slope or threshold values, *p* > .05.

For visual discrimination, a 2 (age) x 2 (reliability) mixed ANOVA of slope values revealed a strong main effect of spatial reliability as expected, *F*(1,43) = 20.186, *p* < .001, *η*^2^ = .32, but no main effect of age or age x reliability interaction. A similar mixed ANOVA of threshold values revealed no significant main effects of age or reliability, nor any interaction, *p* > .05.

No participants were excluded based on their performance in this task. The lowest-performing participant had a fitted slope (accuracy) parameter of 0.10 in the auditory task, which still represents performance well above chance for sounds presented 7° left or right of centre. Auditory threshold (i.e. left/right bias) values were all within 7° (i.e. speaker separation distance) of centre. Similarly, the poorest high reliability visual slope was 0.22, with the most extreme threshold value only 2.52° from centre.

### S2. Motor speed

We used a finger tapping task to compare the age groups in terms of basic motor speed, and to screen participants for significant motor impairment that may affect their ability to respond to the tasks. Participants were instructed to ball their hand into a fist, extending their index finger, and to repeatedly tap a key as quickly as possible for 20 seconds. An on-screen progress bar and countdown provided feedback on performance and time remaining. The task was repeated four times (twice per hand, not including a preceding 10-second practice with each hand).

We analysed the data in terms of the median time between finger taps in seconds (pooled across hands). A two-sample Welch’s *t*-test confirmed that younger participants (*M* = 0.175, *SD* = 0.014) were significantly faster than their older counterparts (*M* = 0.189, *SD* = 0.024), *t*(43) = 2.459, *p* = .018, *d* = 0.73. No participants were excluded based on their performance here, as the slowest responders were within two standard deviations of their respective group means (a conservative threshold).

### S3. GLM analysis of unspeeded ventriloquist paradigm

As well as fitting the Bayesian Causal Inference model, we also performed classical GLM-based analyses on final responses and response times in the unspeeded ventriloquist paradigm. The results are summarised in Figure 2 of the main manuscript.

For auditory localisation responses, the magnitude of the ventriloquist effect was calculated as *VE* = (*A*_*resp*_ − *A*_*loc*_)/(*V*_*loc*_ − *A*_*loc*_) and the mean for each condition was entered into a 2 (age) x 4 (disparity [pooled over direction]) x 3 (visual reliability) mixed ANOVA. This revealed significant main effects of disparity, *F*(3, 129) = 85.31, *p* < .001, *η*^2^ = .67, reliability, *F*(2, 86) = 49.02, *p* < .001, *η*^2^ = .53, and a disparity*reliability interaction, *F*(6, 258) = 30.00, *p* < .001, *η*^2^ = .41. However, no main effect of, or interaction with, age was apparent, *p* > .05. See Figure 2A of the main manuscript.

For common-source judgement responses, we fitted three-parameter Gaussians (peak, mean, standard deviation) to the probability of perceiving a common source as a function of (signed) audiovisual disparity (*A*_*loc*_ − *V*_*loc*_) and compared these parameters separately in 2 (age) x 3 (visual reliability) mixed ANOVAs. The peak of the Gaussian varied with visual reliability level, *F*(2, 84) = 49.24, *p* < .001, *η*^2^ = .54. The mean and width parameters were not significantly affected by visual reliability, and no main effect of (or interaction with) age was apparent for any of the parameters, *p* > .05. See Figure 2C of the main manuscript.

Median response times to the auditory localisation task were analysed in a 2 (age) x 4 (disparity [pooled over direction]) x 3 (visual reliability) mixed ANOVA. Aside from a main effect of age, *F*(1, 43) = 9.77, *p* = .003, *η*^2^ = .19, response times did not differ significantly between conditions, *p* > .05. This is unsurprising, as mouse movements are far more variable (and take much longer) than button presses, so any small effects of condition are likely to be lost. See Figure 2B of the main manuscript.

Median response times to the common-source judgement task were analysed using a 2 (age) x 9 (signed disparity) x 3 (visual reliability) mixed ANOVA. A main effect of age confirmed that older adults were slower overall, *F*(1, 43) = 44.19, *p* < .001, *η*^2^ = .51. Furthermore, age interacted significantly with visual reliability, *F*(2, 86) = 4.92, *p* = .009, *η*^2^ = .09, and audiovisual disparity, *F*(8, 344) = 3.07, *p* = .002, *η*^2^ = .06, and the three-way interaction was also significant, *F*(16, 688) = 2.63, *p* < .001, *η*^2^ = .05. Main effects of reliability, *F*(2, 86) = 9.41, *p* < .001, *η*^2^ = .16, and disparity, *F*(8, 344) = 8.05, *p* < .001, *η*^2^ = .15, were also apparent, as was the interaction between these, *F*(16, 688) = 3.55, *p* < .001, *η*^2^ = .05. See Figure 2C of the main manuscript.

### S4. GLM analysis of speeded ventriloquist paradigm

We performed GLM-based analyses on response times and choices to supplement the compatibility bias model in the speeded ventriloquist task. For these analyses, trials were pooled over left and right locations and hence characterised as spatially congruent or incongruent. Accuracy was quantified as the proportion of correct localisation responses per condition; reaction times were per-condition medians within each participant. We performed four separate 2 (age) x 2 (congruence) mixed ANOVAs, analysing accuracy and reaction times for both the respond-auditory and respond-visual tasks. These results are summarised in panels B and C of Figure 3 of the main manuscript.

A mixed ANOVA of response accuracies in the respond-auditory task revealed a main effect of congruence, *F*(1, 43) = 26.46, *p* < .001, *η*^2^ = .38 (congruent > incongruent), but no main effect of age or interaction, *p* > .05. Conversely, in the respond-visual task a main effect of age showed that older adults were significantly more accurate, *F*(1, 43) = 7.28, *p* = .010, *η*^2^ = .15, possibly due to their higher response threshold (as revealed by the compatibility bias model); a main effect of congruence was also present, *F*(1, 43) = 5.01, *p* = .030, *η*^2^ = .10 (congruent > incongruent), but age and congruence did not significantly interact, *p* > .05.

A mixed ANOVA of response times to the respond-auditory task showed, through a main effect of age, that older adults were significantly slower overall, *F*(1, 43) = 45.01, *p* < .001, *η*^2^ = .51. A main effect of congruence was also present, *F*(1, 43) = 60.57, *p* < .001, *η*^2^ = .51 (congruent < incongruent). Importantly, these factors also interacted, *F*(1, 43) = 15.60, *p* < .001, *η*^2^ = .13, corroborating the finding from the unspeeded paradigm that older adults were disproportionately slower when the task was more challenging (i.e. locating a sound in the presence of an incongruent visual distractor). Main effects of age, *F*(1, 43) = 38.52, *p* < .001, *η*^2^ = .47, and congruence, *F*(1, 43) = 38.89, *p* < .001, *η*^2^ = .48, on response times were also revealed for the respond-visual task, but these did not significantly interact, *p* > .05. See Figure 3B and 3C for a summary.

### S5. BCI model selection

In Section 2.4.2 of the main text we describe a Bayesian Causal Inference model with eleven free parameters, which fitted separate sensory noise and spatial prior parameters depending on the trial type (observers were presented with auditory, visual, and audiovisual signals in separate blocks). However, it is also possible that sensory variances are shared across unisensory and audiovisual blocks, rather than independent. In such a case, the visual and auditory variances would depend only on the external sensory signals and noise imposed by peripheral sensory processing (irrespective of context and task), and the estimation of the auditory and visual variances jointly from all data would provide more precise parameter estimates. If sensory variances are *not* shared across unisensory and audiovisual blocks, treating them as though they are would lead to biased estimation. Likewise, the spatial priors may, or may not, depend on stimulus blocks/task context. To formally address these questions we compared the following three models, which differed in whether the sensory variances and spatial prior were allowed to vary across task context.

Model A (i.e. the standard Bayesian inference model with 6 parameters) assumed that sensory variances and priors were equal for unisensory and bisensory blocks. This model thus included six parameters: *p*_*common*_, *σ*_*P*_, *σ*_*A*_, *σ*_*V1*_, *σ*_*V2*_, *σ*_*V3*_.

Model B constrained the spatial prior to be equal for unisensory and audiovisual blocks, but allowed the sensory variances to differ between unisensory and audiovisual contexts, yielding ten parameters: *p*_*common*_, *σ*_*P*_, *σ*_*A uni*_, *σ*_*V1 uni*_, *σ*_*V2 uni*_, *σ*_*V3 uni*_, *σ*_*A bi*_, *σ*_*V1 bi*_, *σ*_*V2 bi*_, *σ*_*V3 bi*_ (with the indices *uni* and *bi* referring to unisensory and bisensory blocks respectively).

Model C allowed the sensory variances and spatial prior variances to differ between unisensory and audiovisual contexts. Hence, this model included 11 parameters: *p*_*common*_, *σ*_*P uni*_, *σ*_*A uni*_, *σ*_*V1 uni*_, *σ*_*V2 uni*_, *σ*_*V3 uni*_, *σ*_*P bi*_, *σ*_*A bi*_, *σ*_*V1 bi*_, *σ*_*V2 bi*_, *σ*_*V3 bi*_.

We arbitrated between these three models using the Bayesian information criterion (BIC) as an approximation to the model evidence. We performed Bayesian model comparison (Rigoux et al., 2014) at the group (random effects) level as implemented in SPM12 (Stephan et al., 2009; Friston et al., 1994), pooled across age groups, to obtain the protected exceedance probability (the probability that a given model is more likely than any other model, beyond differences due to chance) for the candidate models.

Model C, that fitted sensory variance and spatial prior parameters separately for unisensory and bisensory contexts, outperformed the others at the group level with a protected exceedance probability of 0.58 (compared with values of 0.18 for Model A and 0.24 for Model B). This suggests that the task and stimulus context influenced the estimates of sensory variances and spatial priors to some degree. We therefore report, and compare between groups, the parameters obtained from Model C.

### S6. BCI analysis including motor noise

To account for the possibility of age differences in ability to use a mouse, we fitted a version of our winning BCI model that included *σ*_*motor*_ as an extra free parameter. A summary of the results is given below. It basically replicates the main findings from our main analysis and reveals no significant differences between age groups for motor noise.

**Table S1.**
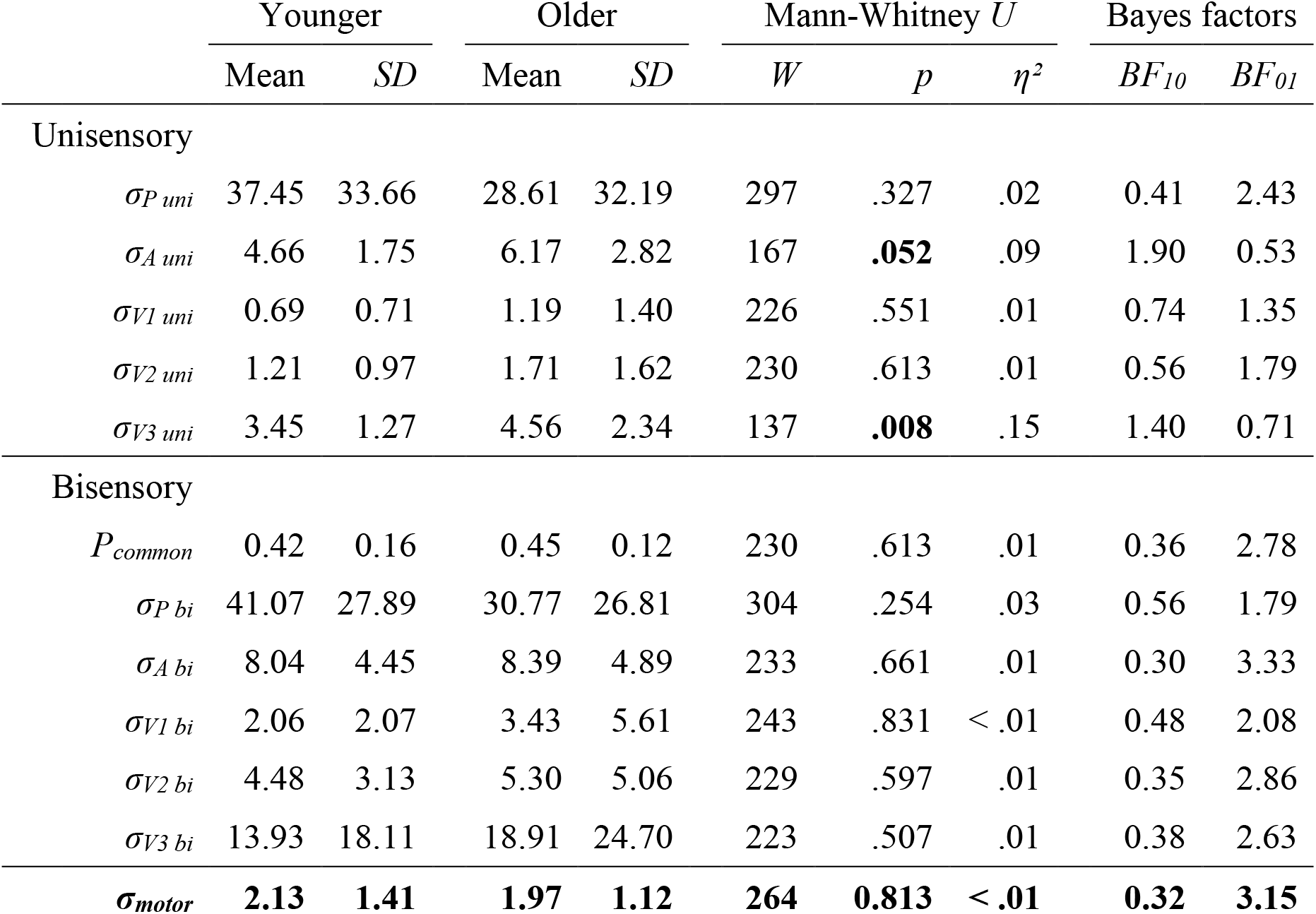
Bayesian Causal Inference parameters (across-participants mean, SD) for younger (n = 23) and older (n = 22) participants, including fitted motor kernel. Mann-Whitney *U* tests with Bayes factors comparing the BCI parameters between older and younger adults. The Bayesian Causal Inference model was fitted jointly to unisensory and audiovisual conditions allowing for separate parameters for the standard deviation of the spatial prior (*σ*_*P,uni*_, *σ*_*P,bi*_) and sensory noise (*σ*_*A,uni*_, *σ*_*A,bi*_, *σ*_*V1 uni*_, *σ*_*V1 bi*_ …). The standard deviation of motor response *σ*_*motor*_ was constrained to be the same for unisensory and multisensory localisation responses. *BF*_10_ quantifies degree of support for the alternative hypothesis that the groups differ, relative to the null hypothesis; *BF*_01_ shows degree of support for the null hypothesis that there is no difference between groups, relative to the alternative hypothesis.

| | Younger | | Older | | Mann-Whitney $U$ | | | Bayes factors | |
| --- | --- | --- | --- | --- | --- | --- | --- | --- | --- |
| | Mean | SD | Mean | SD | $W$ | $p$ | $\eta^2$ | $BF_{10}$ | $BF_{01}$ |
| Unisensory |  |  |  |  |  |  |  |  |  |
| $\sigma_{P\ uni}$ | 37.45 | 33.66 | 28.61 | 32.19 | 297 | .327 | .02 | 0.41 | 2.43 |
| $\sigma_{A\ uni}$ | 4.66 | 1.75 | 6.17 | 2.82 | 167 | **.052** | .09 | 1.90 | 0.53 |
| $\sigma_{V1\ uni}$ | 0.69 | 0.71 | 1.19 | 1.40 | 226 | .551 | .01 | 0.74 | 1.35 |
| $\sigma_{V2\ uni}$ | 1.21 | 0.97 | 1.71 | 1.62 | 230 | .613 | .01 | 0.56 | 1.79 |
| $\sigma_{V3\ uni}$ | 3.45 | 1.27 | 4.56 | 2.34 | 137 | **.008** | .15 | 1.40 | 0.71 |
| Bisensory |  |  |  |  |  |  |  |  |  |
| $P_{common}$ | 0.42 | 0.16 | 0.45 | 0.12 | 230 | .613 | .01 | 0.36 | 2.78 |
| $\sigma_{P\ bi}$ | 41.07 | 27.89 | 30.77 | 26.81 | 304 | .254 | .03 | 0.56 | 1.79 |
| $\sigma_{A\ bi}$ | 8.04 | 4.45 | 8.39 | 4.89 | 233 | .661 | .01 | 0.30 | 3.33 |
| $\sigma_{V1\ bi}$ | 2.06 | 2.07 | 3.43 | 5.61 | 243 | .831 | < .01 | 0.48 | 2.08 |
| $\sigma_{V2\ bi}$ | 4.48 | 3.13 | 5.30 | 5.06 | 229 | .597 | .01 | 0.35 | 2.86 |
| $\sigma_{V3\ bi}$ | 13.93 | 18.11 | 18.91 | 24.70 | 223 | .507 | .01 | 0.38 | 2.63 |
| **$\sigma_{motor}$** | **2.13** | **1.41** | **1.97** | **1.12** | **264** | **0.813** | **< .01** | **0.32** | **3.15** |

